# VDAC activation inhibits hippocampal plasticity via NLRP3 inflammasome & caspase-1: modulation by allopregnanolone enantiomers

**DOI:** 10.1101/2025.10.24.684242

**Authors:** Yukitoshi Izumi, Kazuko A. O’Dell, Douglas F. Covey, Mingxing Qian, Steven Mennerick, Charles F. Zorumski

**Affiliations:** Department of Psychiatry, Washington University in St. Louis School of Medicine St. Louis MO; Taylor Family Institute for Innovative Psychiatric Research, Washington University in St. Louis School of Medicine St. Louis MO; Department of Developmental Biology, Washington University in St. Louis School of Medicine St. Louis MO

**Keywords:** Long-term potentiation, mitochondria, learning, neurosteroids, erastin, metaplasticity

## Abstract

Voltage-dependent anion channels (VDACs) are the most abundant proteins in the outer mitochondrial membrane (OMM) and key regulators of mitochondrial function under physiological and pathological conditions. These channels are modulated by multiple agents and are known binding sites for neuroactive steroids (NAS) including allopregnanolone (AlloP). Using erastin, an agent that promotes VDAC activation by preventing inhibition by tubulin, we assessed the impact of VDAC activation on hippocampal function. Brief erastin administration had no effect on basal transmission but completely inhibited induction of long-term potentiation (LTP) in the Schaffer collateral pathway of rat hippocampal slices. This LTP inhibition was prevented by VBIT-4, an agent that inhibits VDAC oligomerization. VDAC-mediated LTP inhibition was also prevented by inhibitors of the NLRP3 inflammasome and caspase-1, downstream effectors of VDACs, but not by inhibition of cGAS-STING, narrowing the neuroinflammatory pathways involved. Similarly, effects of erastin were prevented by AlloP and at lower concentrations by its unnatural enantiomer (*ent*-AlloP), an agent that unlike AlloP has little effect on GABA_A_ receptors. Erastin also inhibited memory formation in a hippocampal-dependent form of one-trial learning and these effects were prevented by VBIT-4 and *ent*-AlloP, but not by a non-sedating dose of AlloP. These results have relevance for understanding the role of VDACs as mediators of neuronal stress and for the further development of NAS as neurotherapeutics and modulators of cellular stress.

**SIGNIFICANCE STATEMENT:** Voltage-dependent anion channels (VDACs) are important regulators of mitochondrial function, playing roles in cellular metabolism, stress responses and neuroinflammation, and contribute to the pathogenesis of neuropsychiatric illnesses. Here we show that erastin, an agent that activates VDACs initiates specific neuroinflammatory responses to acutely disrupt synaptic plasticity in the rodent hippocampus and abrogate learning. These adverse effects are prevented by the neuroactive steroids allopregnanolone, which binds VDACs and is used clinically for postpartum depression, and its unnatural enantiomer, suggesting that these agents could have therapeutic effects in a broad range of brain illnesses.

## INTRODUCTION

Endogenous neurosteroids that enhance inhibition by potentiating γ-aminobutyric acid type A receptors (GABA_A_Rs), contribute to the pathophysiology of several neuropsychiatric illnesses including postpartum depression (PPD), major depressive disorder (MDD), and anxiety disorders, among others (Zorumski et al., 2013; 2019). These steroids are important stress-induced modulators in the brain and are synthesized in a variety of cell types including excitatory (glutamatergic) neurons (Belelli et al., 2020; 2021). Acute stressors increase these GABAergic steroids, notably allopregnanolone (AlloP) (Purdy et al., 1991), while chronic stress dampens AlloP levels (Antonoudiou et al., 2022; Walton et al., 2023). Human studies have found decreased levels of AlloP in depression and stress-related illnesses (Schule et al., 2014; Maguire and Mennerick, 2024; Meltzer-Brody et al, 2024).

Synthetic neuroactive steroids (NAS) have potential as neurotherapeutics as evidenced by approval of brexanolone and zuranolone for PPD (Deligiannidis et al., 2021; Patterson et al., 2024) and ganaxolone for seizures in a neurodevelopmental illness. GABA_A_R positive allosteric modulation (PAM) plays a key role in the actions of these agents (Maguire and Mennerick, 2024), but NAS have effects that extend beyond GABA_A_Rs and interact with intracellular targets. NAS are protective against forms of neurodegeneration (Langmade et al., 2006; Irwin et al., 2014; Ishikawa et al., 2014), modulate neuroinflammation (Balan et al., 2019; 2021; 2022) and stimulate macroautophagy (Liao et al., 2009; Kim et al., 2012; Ishikawa et al., 2021; 2022).

These latter effects are observed with AlloP, a potent and effective GABA_A_R modulator, and its unnatural enantiomer (*ent*-AlloP), a synthetic agent with marginal effects on GABA_A_Rs (Wittmer et al., 1996; Covey et al., 2023). These observations underlie the importance of identifying non-GABA_A_R targets by which AlloP enantiomers modulate brain function.

NAS-derived photoaffinity labels (PALs) have identified both GABAergic and non-GABAergic targets for AlloP (Zorumski et al., 2025). Major non-GABA targets include voltage-dependent anion channels (VDACs), the most abundant proteins expressed in outer mitochondrial membranes (OMM). VDACs regulate critical mitochondrial functions including ion flux across the OMM, release of metabolites (Camara et al., 2017; Shoshan-Barmatz et al., 2017), neurosteroidogenesis (Marriott et al., 2012; Prasad et al., 2015) and neuroinflammation (Balan et al., 2019; 2021l 2022; Izumi et al., 2024; 2025). VDACs are also involved in pathological processes including release of mitochondrial DNA (mtDNA) and reactive oxygen species (ROS) into cytoplasm (Rostovtseva et al., 2020; Hu et al., 2022). Based on studies with PALs, AlloP analogues bind VDAC1 at a key glutamate residue (E73) in the transmembrane and lipid facing region of the protein (Rister et al., 2023; Darbandi-Tonkabon et al., 2003; 2004). How NAS binding to VDACs influences their function is unknown although AlloP does not alter voltage-dependent channel gating (Budelier et al., 2017; Cheng et al., 2019) and VDACs do not contribute to either GABA_A_R or anesthetic (sedating) effects of AlloP (Darbandi-Tonkabon et al., 2003; 2004).

In the present studies we sought to determine the effects of VDAC activation on synaptic function in the rodent hippocampus and to examine modulation by AlloP enantiomers. For these studies, we used erastin, an anti-tumor agent that activates VDACs by direct binding to the protein (Yagoda et al., 2007) and blocking inhibitory effects of endogenous tubulin on the channels (Shoshon-Barmatz et al., 2017; Rostovtseva and Bezrukof, 2012; Magri et al., 2018). We found that erastin does not alter baseline synaptic function in the hippocampus but inhibits long-term potentiation (LTP), a form of persisting synaptic enhancement that contributes to learning and memory. Effects of erastin are prevented by an agent that inhibits VDAC oligomerization and involve activation of the nucleotide-binding oligomerization domain, leucine-rich repeat and pyrin domain containing 3 (NLRP3) inflammasome and caspase-1 signaling. AlloP enantiomers also prevent the effects of erastin but show reverse enantioselectivity with *ent*-AlloP being more potent than its natural counterpart in both *ex vivo* and *in vivo* experiments.

## METHODS

### Animal Use and Ethics Statement

Sprague-Dawley albino rats were obtained from Harlan Laboratories (Indianapolis IN) and housed under the care of the Washington University Division of Comparative Medicine. All animal use was in accordance with NIH and ARRIVE guidelines. Experimental protocols were approved by the Washington University Institutional Animal Care and Use Committee (IACUC: protocols 22-0220, 22-0228 and 22-0344) and were designed to minimize stress and discomfort.

### Hippocampal slice preparation

Under isoflurane anesthesia and previously published methods (Tokuda et al., 2010; 2011), hippocampi were rapidly dissected from postnatal day (P) 30-32 Harlan Sprague-Dawley male albino rats (Indianapolis IN). Isolated hippocampi were pinned on an agar base in ice-cold artificial cerebrospinal fluid (ACSF) containing (in mM): 124 NaCl, 5 KCl, 2 MgSO_4_, 2 CaCl_2_, 1.25 NaH_2_PO_4_, 22 NaHCO_3_, and 10 glucose gassed with carbogen. A rotary slicer was used to cut the dorsal two-thirds of the hippocampus into 500 µm slices. Slices were then maintained in ACSF at 30°C for at least 1 hour recovery before experiments.

### Hippocampal slice physiology

Single slices were recorded in a submersion-recording chamber with perfusion of ACSF at 30°C and flow rate of 2 ml/min. Extracellular recordings of evoked excitatory postsynaptic potentials (EPSPs) were obtained from the CA1 *stratum radiatum* with 0.1 ms constant current pulses from a bipolar stimulating in the Schaffer collateral (SC) pathway. Stimulus intensity was set at half-maximal based on control input-output (IO) curves. After establishing stable baseline EPSPs, LTP was induced using a single 100 Hz x 1 s high frequency stimulus (HFS) at half-maximal intensity. IO curves were repeated 60 min following tetanic stimulation and were used as the primary measure of synaptic change compared to baseline. Drug concentrations used in these experiments were selected based on having no significant effects on baseline transmission in published reports and our experience with the agents (Izumi et al., 2023).

### Behavioral Studies

To test memory acquisition, we used a one-trial inhibitory avoidance learning task in P30 rats that has previously been shown to involve CA1 hippocampal LTP (Whitlock et al., 2006; Tokuda et al., 2010; Izumi and Zorumski, 2020; Izumi et al., 2021; 2023). In these experiments, the testing apparatus consisted of two chambers one of which was kept constantly lit while the second chamber was maintained in the dark. Both compartments have a floor of stainless-steel rods (4 mm diameter, spaced 10 mm apart) through which an electrical shock can be delivered in the dark chamber. The lit compartment (30 x 20 x 16 cm) was illuminated by four 13 W lights that provided a light intensity of 1000 lux; light intensity in the dark chamber was < 10 lux. On the day of training, animals received a single injection of vehicle (DMSO), VBIT-4 (25 mg/kg ip), AlloP (3 mg/kg ip) or *ent*-AlloP (3 mg/kg) one hour prior to erastin (1 mg/kg ip). One hour after erastin or vehicle, rats were placed in the lit chamber of the testing apparatus and allowed to explore the device for up to 300 s. Immediately after fully entering the dark chamber, a single foot shock was administered and the animal was removed from the apparatus and returned to its home cage. On the next day (24 h after conditioning), rats were placed in the lit chamber without any drug treatment and the latency to enter the dark compartment was recorded over a 300 s trial. The latency to enter the dark chamber was the primary outcome measure. Doses of drugs were determined based on the literature and preliminary experiments and were based on producing minimal sedation over the time course of the conditioning exposure.

### Chemicals

AlloP was obtained from Steraloids, Newport RI (CAS#516-54-1 for physiology studies, Cat#P3810-000) and from Spectrum Chemical (Gardena CA, Cat#A3587) for behavioral studies. We purchased erastin (CAS#571203-78-6 from Selleckchem (Houston TX) for physiology studies (Cat#S7242) and from APExBIO (Boston MA; Cat#B1524 for behavioral studies. VX765 (CAS 273404-37-8, Cat# 7143) was from Tocris (Ellisville MO). MCC950 sodium (CAS 256373-96-3, Cat# S7809) was purchased from SelleckChem (Houston TX). Other chemicals, including LPS (Cat# L2630) and salts, were obtained from Millipore Sigma Chemical Company (St. Louis MO). The AlloP enantiomer was synthesized in the Covey laboratory as described previously (Wittmer et al., 1996; Covey et al., 2023).

### Data Collection & Analysis

Physiological studies were performed and analyzed using pClamp software (Molecular Devices, Union City CA). Results in the text are expressed as mean ± SEM and results from synaptic physiology studies are based on analysis of the maximal rising phase of EPSPs based on IO curves obtained at baseline and 60 min following HFS. EPSPs were normalized to averaged baseline recordings (taken as 100%) and statistical comparisons were performed on responses at 50% maximum of IO curves at baseline and 60 minutes following HFS. A two-tailed unpaired Student’s t-test was used for most comparisons between groups and p < 0.05 was considered significant. When appropriate, paired t-tests were used. For non-normally distributed data, the Mann-Whitney U-test was used for independent samples and the Wilcoxon signed-rank test was used for matched/dependent samples. Numbers reported in the text are the number (N) of animals studied in a condition. Statistics were performed using commercial software (SigmaStat, Systat Software, Inc., Richmond City, CA). Data that are displayed in figures show continuous monitoring of responses at low frequency throughout an experiment and thus may appear to differ from numerical results described in the text, which are based on analysis of IO curves. Behavioral data were analyzed by one-way analysis of variance (ANOVA) followed by Dunnet’s multiple comparison test (Izumi et al., 2023; 2024; 2025).

## RESULTS

### Erastin acutely inhibits hippocampal LTP by activating VDACs

In initial experiments, we examined the effects of acute VDAC activation on synaptic transmission and synaptic plasticity in the CA1 region of rat hippocampal slices using erastin, an agent that activates VDACs by antagonizing inhibition by free tubulin (Yagoda et al., 2007). At a concentration of 0.1 μM, 15-minute perfusion of erastin had no clear effect on basal synaptic transmission but completely inhibited induction of LTP induced by a single 100 Hz x 1 s HFS (Control LTP: 145.2 ± 7.1% change in EPSP slopes measured 60 min following HFS (N=6) vs.87.8 ± 4.7% change 60 min following HFS in the presence of erastin, N=5; p=0. 0001 vs. control LTP; Figure 1A,B).

LTP inhibition by erastin was completely prevented by 2 μM VBIT-4 (Ben-Hail et al., 2016), an agent that inhibits VDAC oligomerization (139.7 ± 11.8% of baseline, N=5; p=0.0035 vs. erastin alone; Figure 1C). When administered alone, VBIT-4 had no effect on LTP induction (138.5 ± 8.6% of baseline, N=6; p=0.5614 vs. control LTP; Extended Figure 1-1A). In contrast to effects on erastin, VBIT-4 had no effect on LTP inhibition by pro-inflammatory stimulation with the toll-like receptor 4 (TLR4) agonist, lipopolysaccharide (LPS) (96.9 ± 9.3%, N=5; Extended Figure 1-1B). However, similar to what we observed with 0.1 μM erastin, we found that 1 μM erastin also had no effect on baseline CA1 transmission but inhibited LTP (96.1 ± 2.4%, N=5; p=0.0002 vs. control LTP; Extended Figure 1-1C) and this LTP inhibition was also largely prevented by VBIT-4 (123.6 ± 1.5%, N=5; p=0.0001 vs. erastin alone; Extended Figure 1-1D). Taken together these results indicate that VDAC activation by acute exposure to erastin profoundly disrupts LTP in the CA1 region.

**Figure 1.**
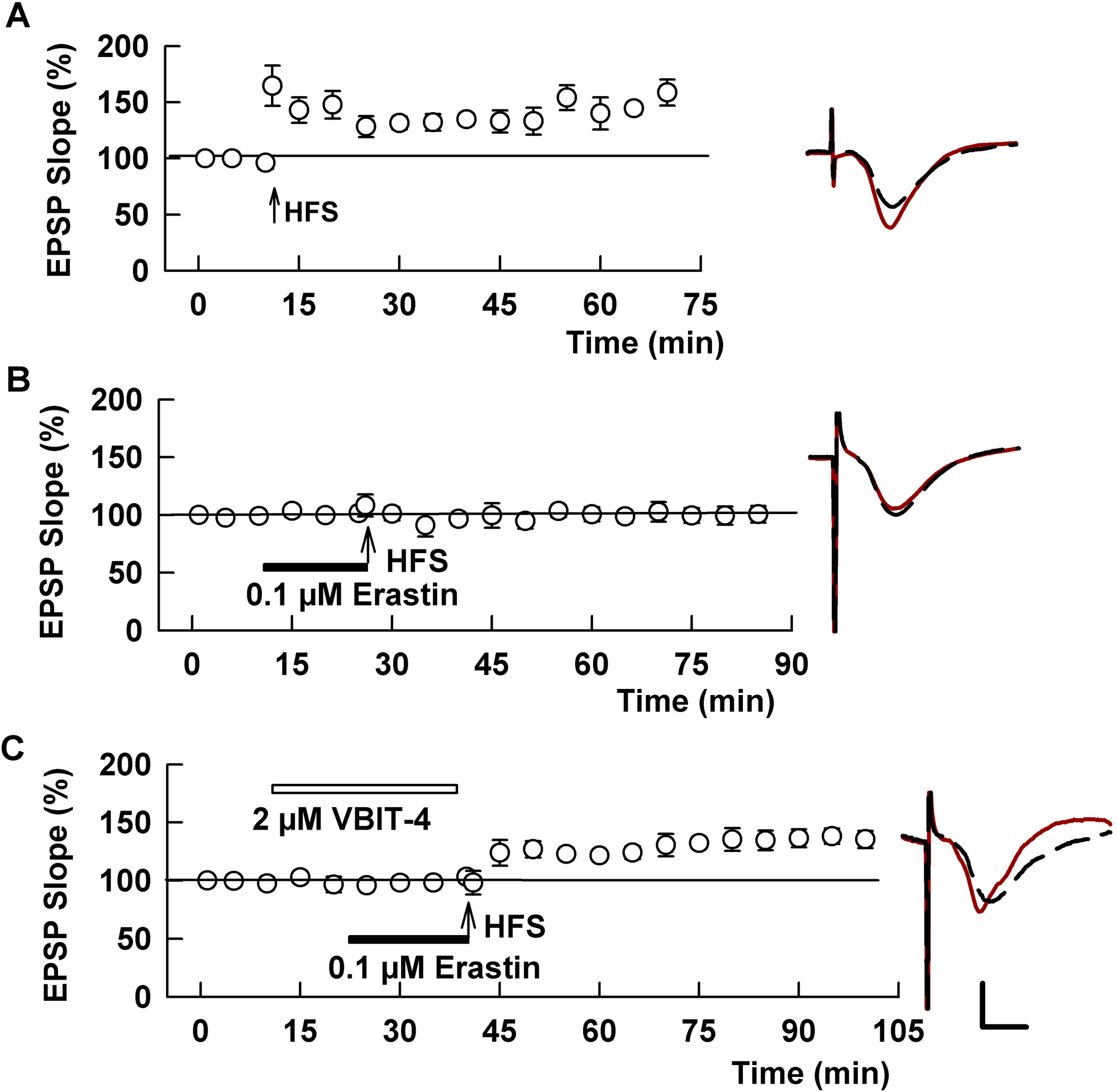
VDAC activation inhibits CA1 hippocampal LTP. A. The graph shows control LTP following a single 100 Hz x 1 s high-frequency stimulation (HFS; arrow) administered to the Schaffer collateral pathway. Traces to the right of the graph in this and other figures show representative EPSPs before (dashed traces) and 60 min following HFS (solid traces). Calibration bar: 1 mV, 5 ms. B. Administration of 0.1 μM erastin (black bar), a VDAC activator, for 15 min prior to HFS completely inhibits LTP induction. C. The effects of erastin are blocked by 2 μM VBIT-4 (white bar), which inhibits VDAC oligomerization, administered prior to and during erastin.

### LTP inhibition by erastin does not involve NMDAR-induced metaplasticity

In prior studies, we found that multiple cellular stressors, including pro-inflammatory stimulation with LPS, disrupt LTP by triggering a form of inhibitory metaplasticity that involves untimely activation of N-methyl-D-aspartate receptors (NMDARs) (Zorumski and Izumi, 2012; Izumi et al., 2021). In these cases, the stressor results in LTP inhibition that outlives stressor exposure for an hour or more, and this prolonged LTP inhibition can be prevented by complete block of NMDARs during the period the stressor is administered. To test whether NMDAR-mediated metaplastic LTP inhibition also occurs with erastin, we treated hippocampal slices with 0.1 μM erastin for 15 min followed by washout for 30 min prior to HFS. Akin to other stressors, this erastin exposure completely inhibited LTP (88.6 ± 2.9% change in EPSPs, N=5, p=0.0001 vs. control LTP; Figure 2A). However, unlike LPS and other stressors (Zorumski and Izumi, 2012; Izumi et al., 2021), this LTP inhibition was not prevented by co-administration of a saturating concentration (50 μM) of the NMDAR antagonist, D-2-amino-5-phosphonovalerate (APV) during erastin exposure (96.2 ± 5.1%, N=4, p=0.2137 vs. erastin alone; Figure 2B).

**Figure 2.**
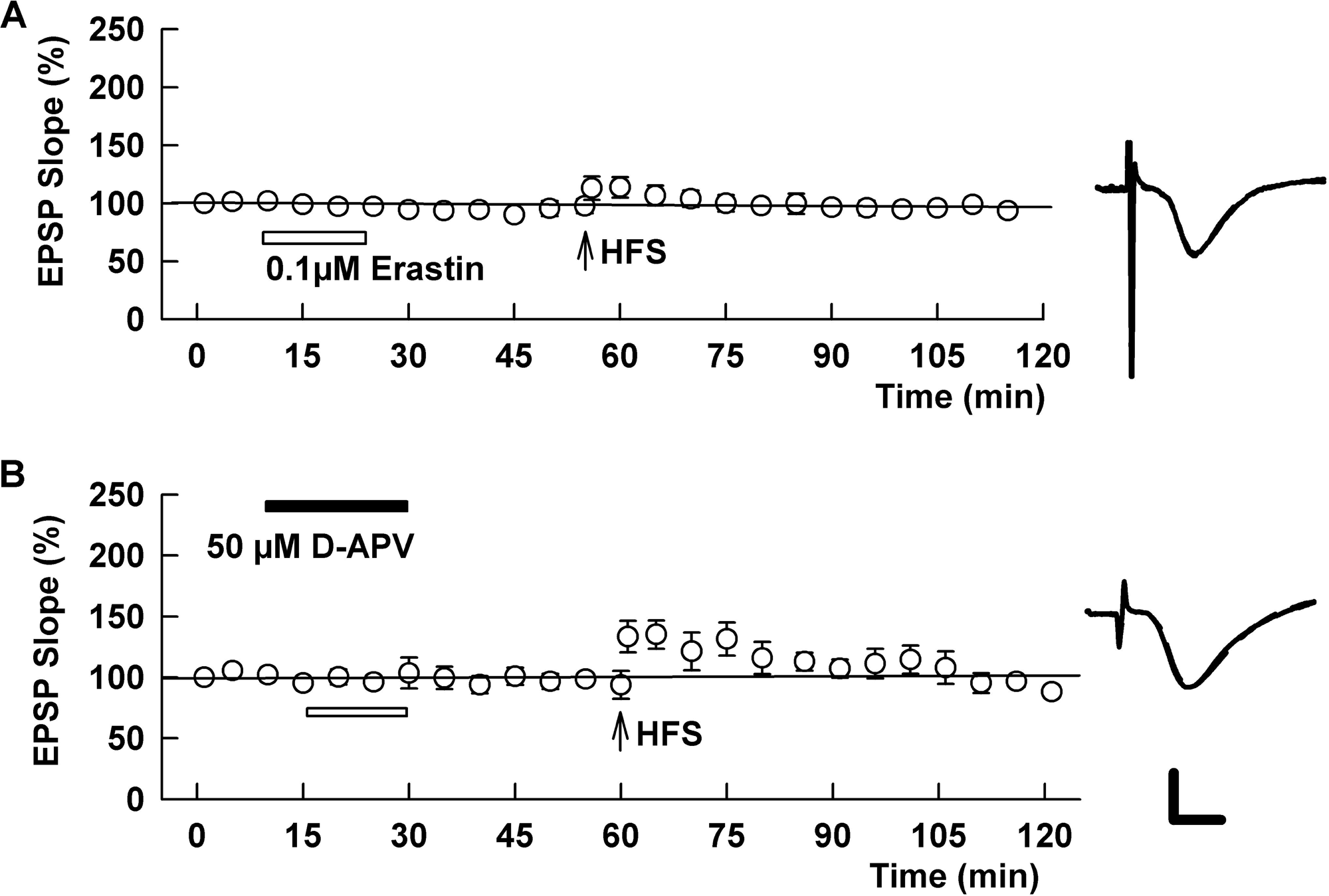
LTP inhibition by erastin persists following drug washout but does not involve metaplastic NMDAR activation. A. Akin to other cellular stressors we have studied (Zorumski and Izumi, 2012), 10 min administration of erastin (white bar) followed by 30 min washout results in persistent LTP inhibition. B. Unlike multiple other stressors (e.g. LPS, Izumi et al. 2021), this persistent LTP inhibition is not altered by co-administration of the competitive NMDAR antagonist, D-APV (black bar). Because D-APV alone inhibits CA1 LTP when administered at the time of HFS, both drugs were washed out for 30 min prior to tetanic stimulation. Traces show representative EPSPs as in Figure 1.

### LTP inhibition by erastin involves the NLRP3 inflammasome but not cGAS-STING

VDAC activation can promote cellular stress and is known to trigger pro-inflammatory signaling including activation of the NLRP3 inflammasome (Zhou et al., 2011; Kelley et al., 2019). Thus, we also examined whether this pathway contributes to effects on synaptic plasticity. Consistent with a role for NLRP3, we found that administration of the NLRP3 inhibitor, MCC950 (Coll et al., 2015) at 0.1 μM prevented erastin-mediated LTP inhibition (EPSP slopes: 137.3 ± 2.8% of baseline, N=5, p=0.0001 vs. erastin alone; Figure 3A). NLRP3 activation occurs upstream of caspase-1 (Bonam et al., 2024) prompting us to examine the effects of VX-765, a small molecule inhibitor of this enzyme (Jin et al., 2022). Similar to MCC950, 1 μM VX765 also completely prevented the acute adverse effects of erastin (150.3 ± 5.7%, N=5, p=0.0001 vs. erastin alone; Figure 3B).

**Figure 3.**
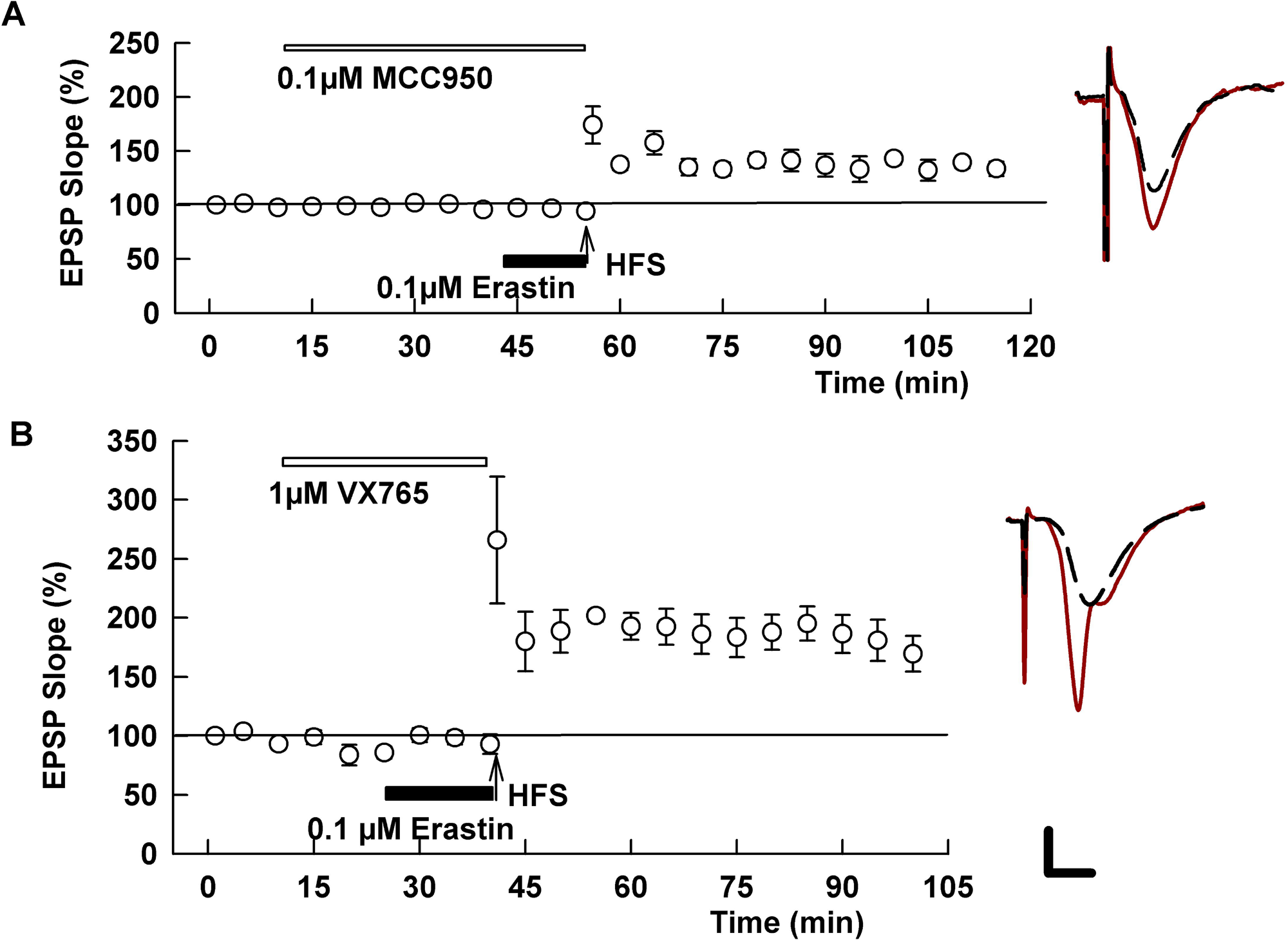
Erastin-mediated LTP inhibition involves activation of the NLRP3 inflammasome and caspase-1. A. In the presence of the NLRP3 inhibitor, MCC-950 (white bar), the adverse effects of erastin on LTP are completely prevented. B. Similarly, the caspase-1 inhibitor, VX-765, blocked the effects of erastin. Traces show representative EPSPs.

VDAC activation can also stimulate the cyclic GMP-AMP synthase-stimulator of interferon genes (cGAS-STING) pathway as a mechanism contributing to cellular stress and inflammatory signaling (Kim et al., 2019). To examine a role for cGAS-STING in the effects of erastin we used the STING inhibitor, H-151 (Kim et al., 2019; Kobritz et al., 2022). At a concentration of 1 μM, H-151 alone completely inhibited CA1 LTP (94.2 ± 7.9%; n=5; Extended Figure 3-1A). This result is consistent with a prior report showing that H-151 knockout disrupts hippocampal-dependent learning (Neupane et al., 2024). A lower concentration of H-151 allowed LTP induction but had no effect on erastin-mediated LTP inhibition (0.1 μM H-151 alone: 139.7 ± 11.8%, N=5; 0.1 μM H-151 + erastin: 98.5 ± 8.7%, N=5; p=0.0228 vs. H-151 alone; Extended Figure 3-1B,C).

VDACs, particularly VDAC-2, also promote synthesis of endogenous 5α-reduced neurosteroids such as AlloP (Marriott et al., 2012; Prasad et al., 2015), and AlloP via effects on GABA_A_Rs can adversely modulate LTP induction (Zorumski and Izumi, 2012). To test a role for AlloP in the effects of erastin we used dutasteride, a broad-spectrum inhibitor of the key neurosteroid synthetic enzyme, 5-alpha reductase. At 1 μM, however, dutasteride had no effect on LTP inhibition by erastin (100.6 ± 5.8% of baseline, N=5; Extended Figure 3-1D). Thus, endogenous 5α-reduced neurosteroid production appears to play no role in the erastin effects observed here.

### Activation of macroautophagy prevents the effects of erastin

These results indicate that activation of VDAC by erastin triggers the NLRP3 inflammasome to inhibit hippocampal synaptic plasticity. VDAC has several effects that promote NLRP3 activation including release of reactive oxygen species (ROS) and/or oxidized mitochondrial DNA (mtDNA) into the cytoplasm (Varughese et al., 2021; Zhao et al., 2021). Inflammasome activation can be dampened by the homeostatic cellular mechanism of autophagy, which limits NLRP3 activation by removing activating stimuli (Zhao et al., 2021). Thus, strategies to stimulate autophagy could dampen the impact of VDAC activation. To test this hypothesis, we used 200 nM rapamycin, an agent that activates autophagy by inhibiting mechanistic target of rapamycin (mTOR), a kinase that plays a key role in regulating autophagy (Ishikawa et al., 2021; 2022). In the presence of rapamycin, erastin had no effect on LTP (145.4 ± 16.0%, N=5; p=0.0086 vs. erastin alone; Figure 4B). Similarly, Torin-1, which also stimulates autophagy by inhibiting mTOR (Ishikawa et al., 2021), prevented LTP inhibition when administered at 1 μM prior to and during erastin (140.6 ± 9.4%, N=5; p=0.0010 vs. erastin alone; Figure 4A).

**Figure 4.**
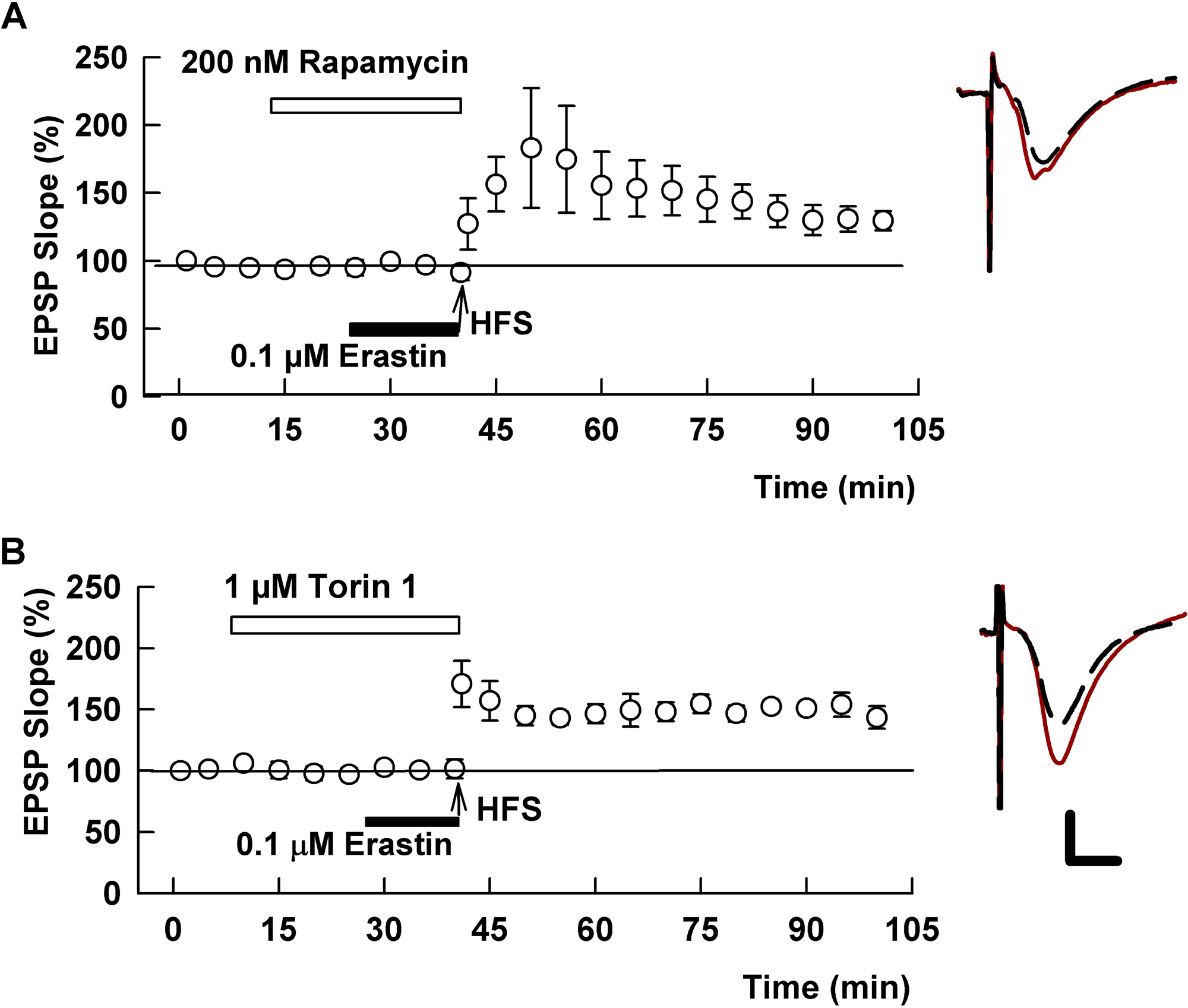
Agents that activate macroautophagy prevent the effect of erastin on LTP. A. In the presence of 200 nM rapamycin, an inhibitor of mTOR that activates autophagy, erastin fails to alter LTP induction. B. Similarly, the adverse effects of erastin are prevented by 1 μM Torin-1, an agent that more specifically inhibits mTOR to promote autophagy. Traces show representative EPSPs.

### AlloP enantiomers modulate the effects of erastin on LTP

The neurosteroid, AlloP, also stimulates autophagy as a neuroprotective mechanism under stressful conditions (Ishikawa et al., 2021). In contrast to the well-known effects of AlloP on GABA_A_Rs that are highly enantioselective (Wittmer et al., 1996; Covey et al., 2023), neuroprotective effects of AlloP are not enantioselective and the GABA-inactive AlloP enantiomer also stimulates autophagy (Ishikawa et al., 2022). Furthermore, AlloP analogues that are photoaffinity labels directly bind VDAC-1 and VDAC-2 but have unknown effects on these proteins (Cheng et al., 2019). These observations prompted us to examine whether AlloP and *ent*-AlloP modulate effects of erastin on hippocampal plasticity. In initial experiments, we found that 1 μM AlloP, a concentration that has anti-inflammatory effects against LPS and other Toll-like receptor agonists (Izumi et al., 2024) had no effect on LTP inhibition by erastin (91.5 ± 6.3%, N=5, Figure 5A). Although this result was observed with a relatively short administration of AlloP, we also found that more prolonged AlloP administration (2-4 h) had no effect on LTP inhibition (102.0 ± 5.6%, N=7, Extended Figure 5-1A). However, in contrast to 1 μM AlloP, a higher concentration of AlloP (5 μM) prevented the effects of erastin using the shorter drug administration (149.4 ± 11.8%, N=4; p=0.0013; Figure 5B).

Because *ent*-AlloP is also neuroprotective and promotes autophagy (Ishikawa et al., 2022), we examined its effect against erastin. In contrast to natural AlloP, a 2-4 h incubation of slices with 1 μM *ent*-AlloP completely prevented LTP inhibition by 0.1 μM erastin (138.8 ± 7.9%, N=5; p=0.0005 vs. erastin alone; Extended Figure 5-1B) and by 1 μM erastin (133.5 ± 15.0%, N=5; p=0.0392 vs. 1 μM erastin; Extended Figure 5-1C). These results prompted us to examine a shorter application of *ent*-AlloP that has been found to have anti-inflammatory effects (Izumi et al.,2024). When administered for 15 min prior to and during 0.1 μM erastin, *ent*-AlloP also prevented LTP inhibition (143.2 ± 4.5%, N=5; p=0.0001 vs. erastin; Figure 5B). Similar results were obtained with 15 min applications of 0.5 μM *ent*-AlloP (163.4 ± 21.0%, N=3) supporting clear differences in potency between the AlloP enantiomers.

**Figure 5.**
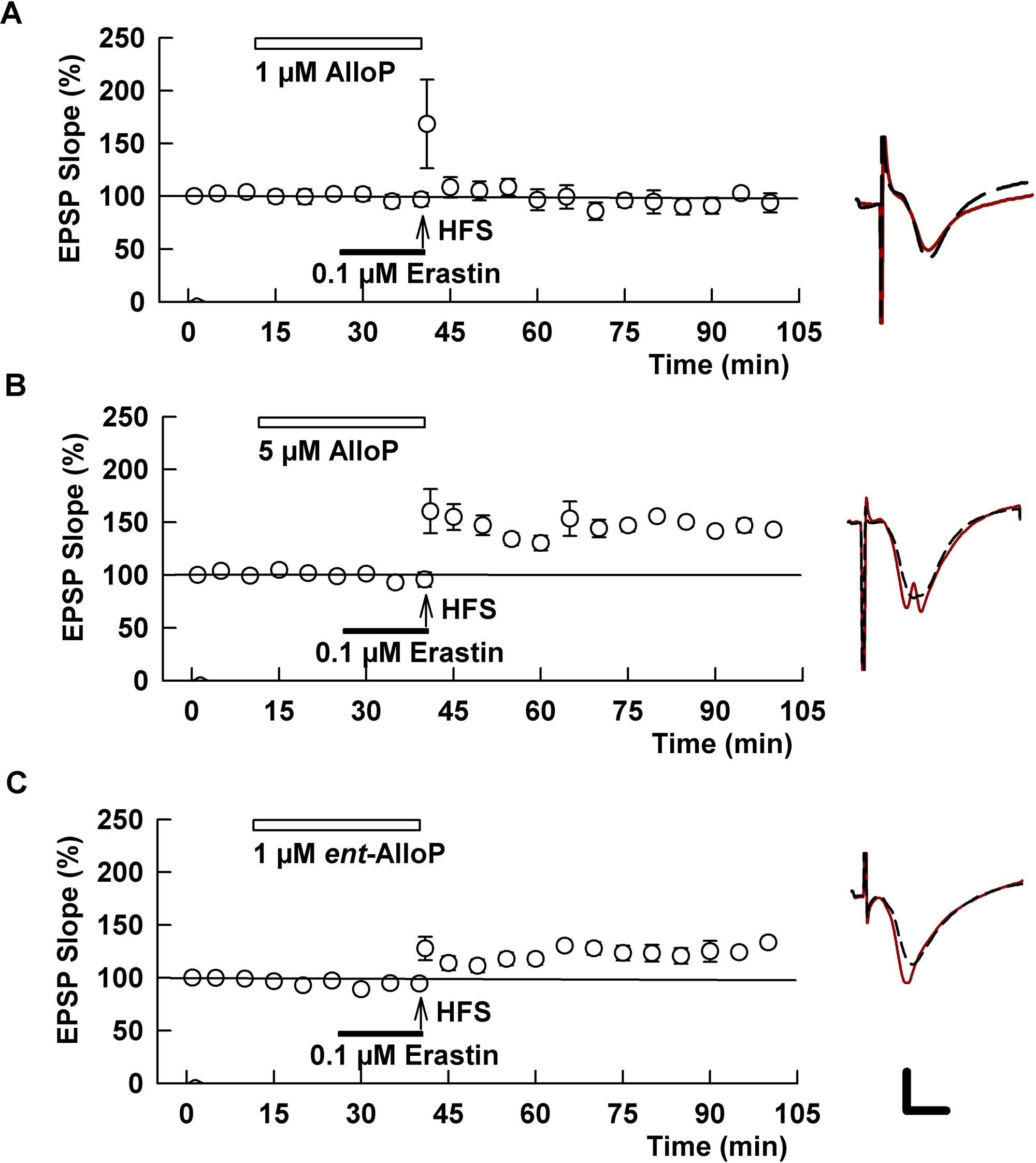
AlloP enantiomers prevent the effects of erastin on CA1 LTP but show reverse enantioselectivity. A. When administered at 1 μM prior to and during erastin, AlloP has no effect on LTP inhibition. B. A higher concentration of AlloP (5 μM) completely prevented erastin-mediated LTP block. C. In contrast to AlloP, the unnatural AlloP enantiomer (*ent*-AlloP) prevented the effects of erastin at 1 μM. Traces show representative EPSPs as in Figure 1.

Based on the protective effects of autophagy activators and neurosteroids described above (Figure 4), we sought to determine whether effects on autophagy contribute to the actions of the AlloP enantiomers using agents that are known to inhibit autophagy. However, when slices were preincubated with 0.1 μM bafilomycin A1, an inhibitor of vesicular ATPase that also impairs loading of glutamate into synaptic vesicles, LTP induction was completely inhibited (94.5 ± 7.9% of baseline, N=5; p=0.0010 vs control LTP); this LTP inhibition was not prevented by either 5 μM AlloP (105.8 ± 9.6%, N=8, p=0.4290 vs bafilomycin alone; Extended Figure 5-2A) or 1 μM *ent*-AlloP (95.6 ± 7.5%, N=5; p=0.9221 vs. bafilomycin, Extended Figure 5-2B). Similarly, 1 μM Autophagy Inhibitor VII (AUIVII, acridinamine 33), an agent that blocks autophagy by inhibiting the Class III phosphoinositide 3-kinase complex (Wong et al., 2015), also disrupted LTP (110.5± 6.4%, N=5; p=0.0061 vs. control LTP) and its effects were not altered by *ent*-AlloP (93.0 ± 4.6%, N=5; p=0.0572 vs. AUVII alone, Extended Figure 5-2C). We also found that a third autophagy inhibitor, 3-methyl adenosine (3MA) blocked LTP on its own (101.3 ± 2.8%, N=5; p=0.0005 vs. control LTP) and its effects were not altered by *ent*-AlloP (94.5 ± 3.9%, N=5; Extended Figure 5-2D). These results are consistent with the important role of basal autophagy in regulating synaptic function and plasticity (Glatigny et al., 2019; Pandey et al., 2021; Karpova et al., 2025) and suggest that the effects of the AlloP enantiomers against erastin likely occur upstream of autophagy activation.

### Erastin impairs hippocampal dependent learning: modulation by AlloP enantiomers

To determine whether results on LTP in hippocampal slices translate to behavioral changes, we examined the effects of erastin, VBIT-4 and the AlloP enantiomers on one-trial inhibitory avoidance learning, a learning task dependent upon hippocampal LTP (Whitlock et al. 2006).

For this task, rats are placed in the lit chamber of a two-chamber apparatus. Once rats enter the dark compartment, they are given a foot shock and returned to their home cage. Twenty-four hours later, animals are tested for memory retention (Figure 6A). Rats treated with vehicle (DMSO) one hour prior to conditioning learn the task well and remain in the lit compartment for the full 300 s test period (Figure 6B). Animals receiving erastin (1 mg/kg ip) one hour prior to conditioning exhibit impaired learning 24 h later (remaining in the lit chamber for only 86.0 ± 32.9 s, N=6, p <0.0001 vs. vehicle controls). The effects of erastin were completely prevented by the VDAC inhibitor, VBIT-4 (25 mg/kg ip) (time remaining in the lit chamber = 300 s, N=5; p < 0.0001 vs. erastin alone). Pretreatment with AlloP (3mg/kg ip) showed variable but non-significant effects on erastin-mediated impairment (remaining time: 160.8 ± 62.7 s, N=6; p = 0.1570). We attempted to use a higher dose of AlloP but that dose caused acute behavioral sedation that precluded accurate conditioning and memory testing. In contrast, *ent*-AlloP (3 mg/kg ip) was non-sedating and prevented the erastin-induced learning impairment (remaining time: 278.6 ± 21.4 s, N=8; p<0.0001). Overall, these behavioral results are consistent with results observed in LTP experiments including differences in apparent potency between the AlloP enantiomers.

**Figure 6.**
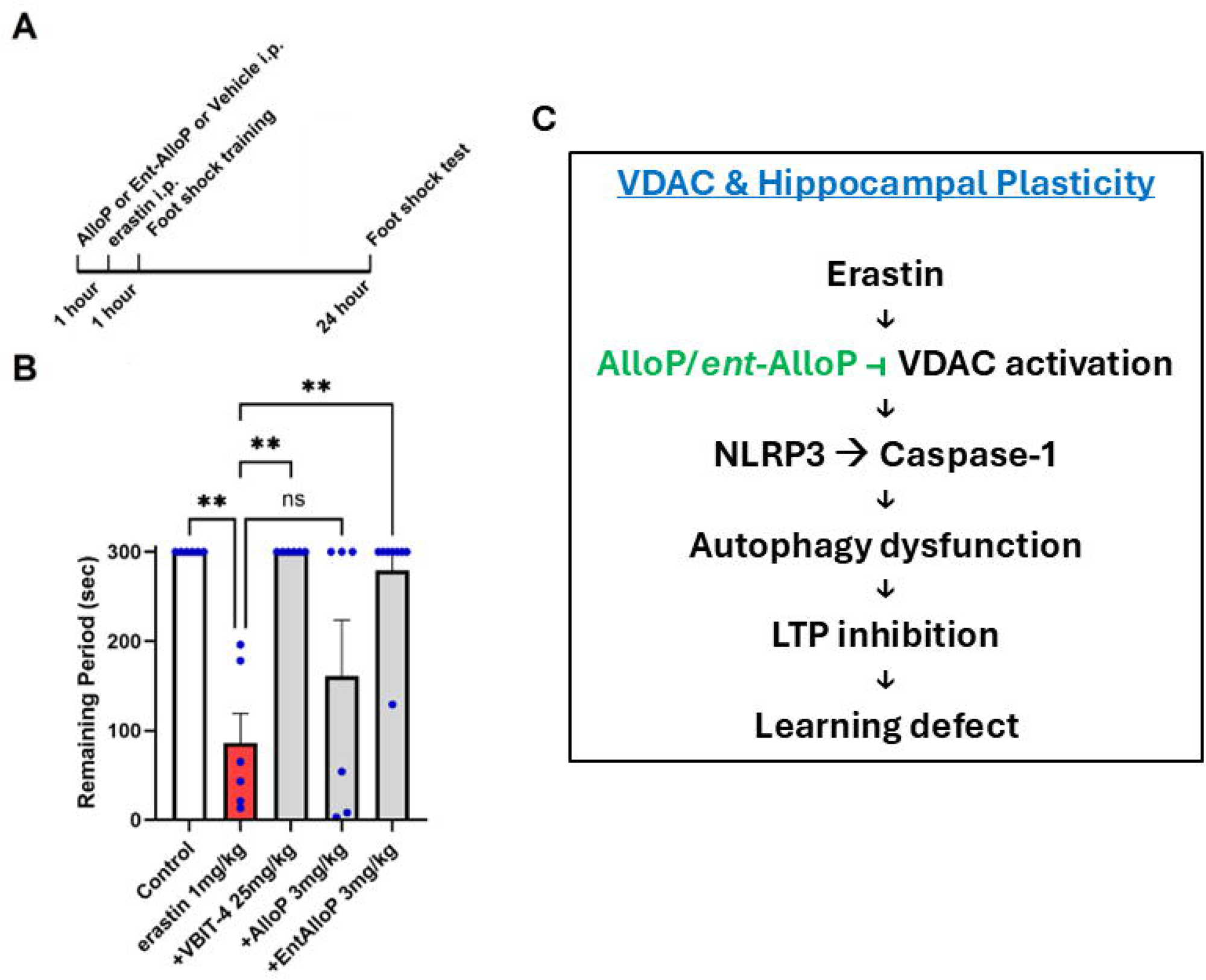
VDAC activation results in defects in one-trial inhibitory avoidance learning. A. The diagram depicts the experimental paradigm used to study the effects of erastin on learning in the absence and presence of the VDAC inhibitor and the AlloP enantiomers. B. When erastin (1 mg/kg ip) was administered 1 h prior to conditioning, animals displayed markedly impaired learning and memory. In contrast, control rats treated with vehicle alone (DMSO) failed to leave the lit chamber during the 5 min testing period 24 hours following conditioning. Both VBIT-4 (25 mg/kg ip) and *ent*-AlloP (3 mg/kg ip) prevented the adverse effects of erastin. However, AlloP (3mg/kg ip) had variable but insignificant effects. Higher doses of AlloP in this paradigm resulted in significant sedation that impaired behavioral testing. C. The diagram depicts a proposed cascade of events involved in the effects of VDAC activation with erastin on hippocampal LTP and learning. In this model, the AlloP enantiomers are proposed to act at VDAC, a known binding site. The cascade is linear in this depiction but beyond VDAC there could be parallel effects on NLRP3 and autophagy.

## DISCUSSION

Our results indicate that acute activation of VDACs by erastin has little effect on baseline transmission at Schaffer collateral synapses but disrupts induction of LTP, a form of plasticity that contributes to learning and memory. In turn, VDAC stimulates the NLRP3 inflammasome-caspase 1 pathway to promote LTP inhibition and learning impairment. A proposed scheme for these results is shown in Figure 6C.

These studies provide support for the role of VDACs as regulators of neuronal physiology and plasticity (Camara et al., 2017; Magri et al., 2018; Varughese et al., 2021; Weeber et al., 2002). The VDAC family includes three proteins, VDAC-1, -2 and -3. Of these VDAC-1 and possibly VDAC-2 are likely contributors to our observations. VDAC-1, the most abundant OMM protein, regulates release of ions, metabolic products, lipids and ROS across the OMM (Shoshan-Barmatz et al., 2017). Under basal conditions, VDACs exist as monomers but under cellular stress form oligomers that alter function. VDACs also have scramblase activity and facilitate lipid entry into mitochondria from the endoplasmic reticulum (ER), an effect not dependent upon channel gating (Jahn et al., 2023). VDAC-1 is a major intracellular hub with over 100 partners that influence its function (Shoshan-Barmatz et al., 2017). VDACs are also components of mitochondrial associated membranes (MAMs) where they interact with mitochondrial and ER proteins to regulate inter-organelle interactions (Barbaro-Camps et al., 2014; Gatliff et al., 2014). This latter role includes cholesterol trafficking from ER to mitochondria for steroidogenesis with VDAC-2 playing a key role (Marriott et al., 2012; Prasad et al., 2015).

VDAC is highly regulated and dimeric α/β-tubulin inhibits channel activity (Shoshan-Barmatz et al., 2017; Kim et al., 2019). Erastin blocks the interaction of tubulin with VDAC and promotes oligomerization and channel activation (Maldonado, 2017; Rostovtseva and Bezrukov, 2019). While erastin has effects other than modulation of VDACs, including inhibition of System Xc^-^ cystine-glutamate transporters (Zhao et al., 2020), our results with VBIT-4, an inhibitor of VDAC oligomerization (Ben-Hail et al., 2016), strongly implicate effects on VDACs as a key mechanism underlying erastin-induced LTP defects. Recent studies indicate that VBIT-4 acts by membrane effects rather than direct VDAC interactions (Ravishankar et al., 2025) and can be toxic at concentrations >10 μM. However, we found no toxic effects of VBIT-4 either on basal transmission or LTP under the conditions used and VBIT-4 prevented adverse effects of erastin. Through effects on VDAC, erastin produces major changes in mitochondrial function, including a metabolic switch from aerobic glycolysis to respiratory metabolism and generation of ROS (Maldonado, 2017). The net effect is an increase in cellular stress that activates the NLRP3 inflammasome and caspase-1 (Bonam et al., 2024; Hu et al., 2022). In previous studies, we also found that activation of NLRP3-caspase-1 plays a key role in the adverse effects of pro-inflammatory stimulation with LPS on hippocampal function (Izumi et al., 2025).

VDACs also bind AlloP and other NAS with at least four sites identified on VDAC1 that also bind cholesterol (Budelier et al., 2017; Cheng et al., 2019). The best characterized NAS site on VDAC1 is E73, a glutamate residue in the outer transmembrane region. E73 faces membrane lipid away from the VDAC channel and interacts with multiple agents that modulate VDAC activity including hexokinase, a key mediator of aerobic glycolysis (Rister et al., 2023).

Cholesterol and NAS bind E73 in opposite orientations (Cheng et al., 2019) and E73 is important for VDAC oligomerization. The effect of AlloP on VDAC is not certain but AlloP does not alter voltage-dependent channel gating (Cheng et al., 2019) and VDACs do not contribute to NAS-mediated modulation of GABA_A_Rs or anesthetic effects (Darbandi-Tonkabon et al., 2003; 2004). It appears likely that neurosteroids modulate interactions of VDACs with other proteins and/or oligomerization (Rostovtseva et al., 2020), perhaps by affecting the protonation state of E73 (Bergdoll et al., 2018).

Our results emphasize key roles of VDACs in regulating cellular function via proinflammatory responses. Activated VDACs promote cytoplasmic release of ROS that stimulate the NLRP3 inflammasome and caspase-1 (Zhou et al., 2011). Release of mitochondrial DNA (mtDNA) via VDACs also activates inflammatory responses, including the cGAS-STING pathway and the senescence associated signaling pathway (SASP) (Kim et al., 2019). NLRP3 activation in turn dampens autophagy, a homeostatic cellular stress response that helps to preserve function by recycling damaged proteins and organelles to generate energy (Kelley et al., 2019; Bonam et al., 2024). Here we found that inhibitors of NLRP3 and caspase-1, but not cGAS-STING, prevent effects of erastin on LTP as did agents that stimulate autophagy. In contrast, we found that inhibitors of autophagy disrupted LTP under basal conditions, supporting the importance of autophagy in maintaining synaptic homeostasis (Lieberman and Sulzer, 2020; Liang, 2019).

The present results highlight complex roles that neurosteroids play in cellular stress. We previously found that endogenous 5α-reduced neurosteroids such as AlloP play a key role in mediating effects of a variety of stressors (Zorumski and Izumi, 2012) including pro-inflammatory stimulation (Izumi et al., 2024, 2025). In these studies, AlloP serves as a downstream effector that diminishes synaptic plasticity by enhancing GABAergic inhibition, a well-known effect of AlloP (Zorumski et al., 2019; 2025). In this role, the neurosteroids appear to function as homeostatic neuroprotectors that dampen synaptic plasticity in the face of neural stress (Izumi et al., 2024; Ishikawa et al., 2014). VDAC activation with erastin differs from these observations and does not involve endogenous 5α-reduced neurosteroids based on experiments with dutasteride, a potent 5-alpha reductase inhibitor. However, AlloP also modulates and dampens neuronal stress when administered prior to and during stress (Zorumski and Izumi, 2012). We observed this stress modulation with stimulation of several Toll-like receptors that participate in neuroinflammation (Izumi et al., 2024). An intriguing feature of this stress modulation is that both enantiomers of AlloP can be effective (Izumi et al., 2024; Ishikawa et al., 2021; 2022). This latter observation is important because *ent*-AlloP unlike AlloP has little effect on GABA_A_Rs (Wittmer et al., 1996; Covey et al., 2001; 2023) while both enantiomers accumulate similarly in plasma membranes and intracellular compartments (Jiang et al., 2016). Here we found that both AlloP enantiomers are effective against erastin, with *ent*-AlloP being more potent than AlloP.

While definitive mechanisms by which AlloP enantiomers protect against erastin remain uncertain, there are several possibilities. First, AlloP enantiomers may inhibit VDAC oligomerization via direct effects on the protein, particularly at VDAC-1 E73 or VDAC-2 E84 to prevent dissociation of hexokinase and NLRP3 activation (Baik et al., 2023; Bergdoll et al., 2018; Rister et al., 2023). Second, both AlloP enantiomers stimulate macroautophagy (Liao et al., 2009; Kim et al., 2012), and *ent*-AlloP appears to be more effective than natural AlloP (Ishikawa et al. 2021; 2022). In the present study, autophagy activators prevented effects of erastin on LTP. Third, neurosteroids prevent pro-inflammatory stimulation by certain Toll-like receptors (TLRs) (Balan et al., 2021; Izumi et al., 2024). Although TLRs are unlikely involved in the effects of erastin, anti-inflammatory effects could occur downstream of VDAC at the level of NLRP3 or caspase-1. Autophagy again could be a contributing mechanism based on bidirectional interactions of inflammasome signaling with autophagy (Zhou et al., 2011; 2019; Biasizzo and Kopitar-Jerala, 2020; Bonam et al., 2024). Alternatively, AlloP enantiomers can activate pregnane X receptors (Langmade et al., 2006) and possibly liver X receptors (Divya et al., 2025) to promote anti-inflammatory and neuroprotective effects. Finally, neurosteroids modulate mitochondrial function with beneficial effects on ATP production (Grimm et al., 2014; Lejri et al., 2017). This is an intriguing possibility that could involve VDACs or other steps in mitochondrial energy production.

These results also have implications for understanding neuropsychiatric disorders and further development of NAS as neurotherapeutics (Belelli et al., 2020; 2121; Irwin et al., 2014). There is increasing interest in the role of mitochondrial dysfunction as a mechanism in multiple illnesses including neurodegenerative disorders such as Alzheimer’s disease (Soumalainen and Nunnari, 2024; Argueti-Ostrovsky et al., 2025). Similarly, mitochondrial dysfunction may contribute to primary psychiatric illnesses including mood, anxiety and psychotic disorders (Manji et al., 2012; Pei and Wallace, 2018). Mitochondrial abnormalities include changes in the expression and function of VDACs (Scaini et al., 2019; Lorenzo et al., 2023; Segev et al., 2023). Psychiatric illnesses are also recognized as major contributors to premature aging, adding to illness-associated morbidity (Lorenzo et al., 2023; Wertz et al., 2021). Inflammation is an important mechanism contributing to cellular senescence and aging, and VDACs play important roles in these processes (Kim et al., 2019; Li et al., 2024). Thus, beneficial effects of NAS on VDAC function (Zhou et al., 2019) or OMM permeability (Sayeed et al., 2009) could have therapeutic implications for dampening adverse outcomes in psychiatric illnesses. The effects of *ent*-AlloP on inflammation (Izumi et al., 2024), autophagy (Ishikawa et al., 2022) and VDAC-induced mitochondrial dysfunction in the absence of GABAergic effects may provide a pathway to develop novel treatments devoid of sedation and possible abuse/misuse potential.

## Conflict of Interest

The work was funded by NIMH grants MH123748 (SM) & MH122379 (CFZ, SM), the Taylor Family Institute for Innovative Psychiatric Research (DFC, SM, CFZ) and the Bantly Foundation (CFZ). A

## Acknowledgements

The authors thank members of the Taylor Family Institute for Innovative Psychiatric Research and the Silvio O. Conte Neuroscience Center at Washington University for discussion and input, particularly Alex S. Evers and Jamie Maguire.

## Author contributions

Conceptual study design: CFZ, YI & SM. Data collection and analysis: YI, KO & CFZ. Critical Resources: DFC, CM. Original draft: CFZ. Critical revisions: YI, KO, DFC, CFZ, & SM.

## Notes

### Competing Interest Statement

CFZ previously served as a member of the Scientific Advisory Board for Sage Therapeutics and held equity in the company. Sage Therapeutics had no role in the design or interpretation of these experiments. The remaining authors declare no competing financial interests.

