## Supplementary figures and images for "VDAC activation inhibits hippocampal plasticity via NLRP3 inflammasome & caspase-1: modulation by allopregnanolone enantiomers"

### Extended Figure 1-1

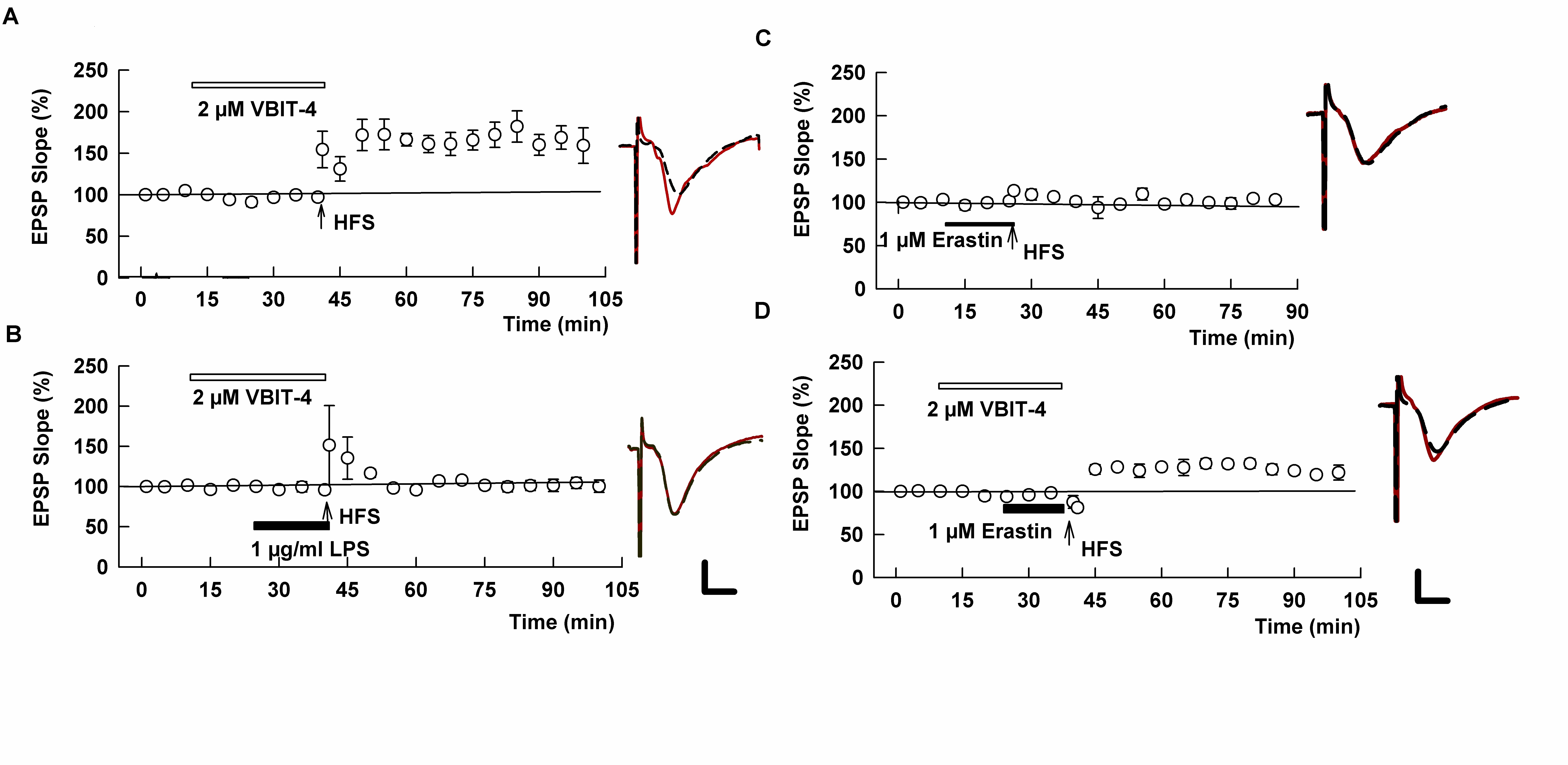

### Extended Figure 3-1

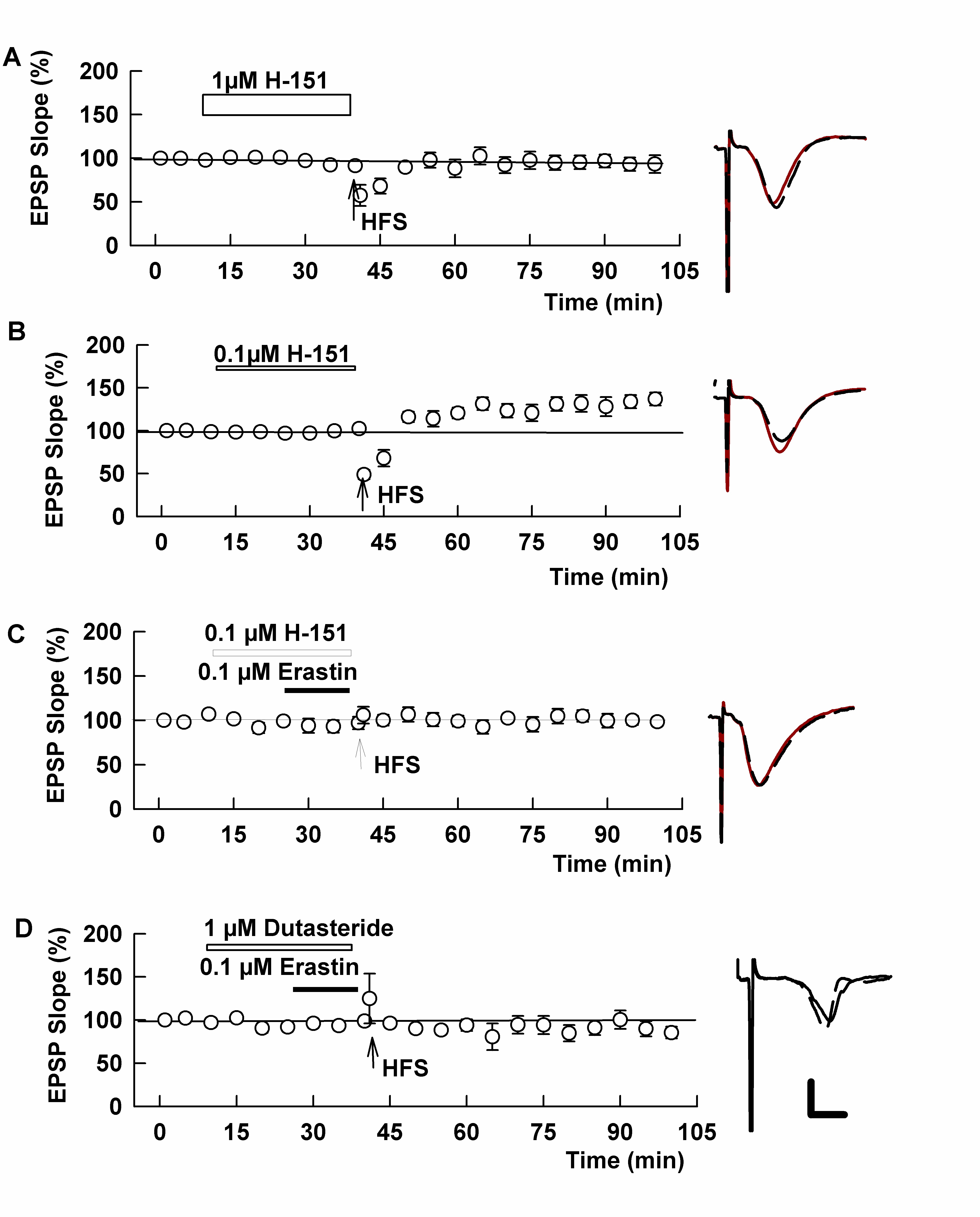

### Extended Figure 5-1

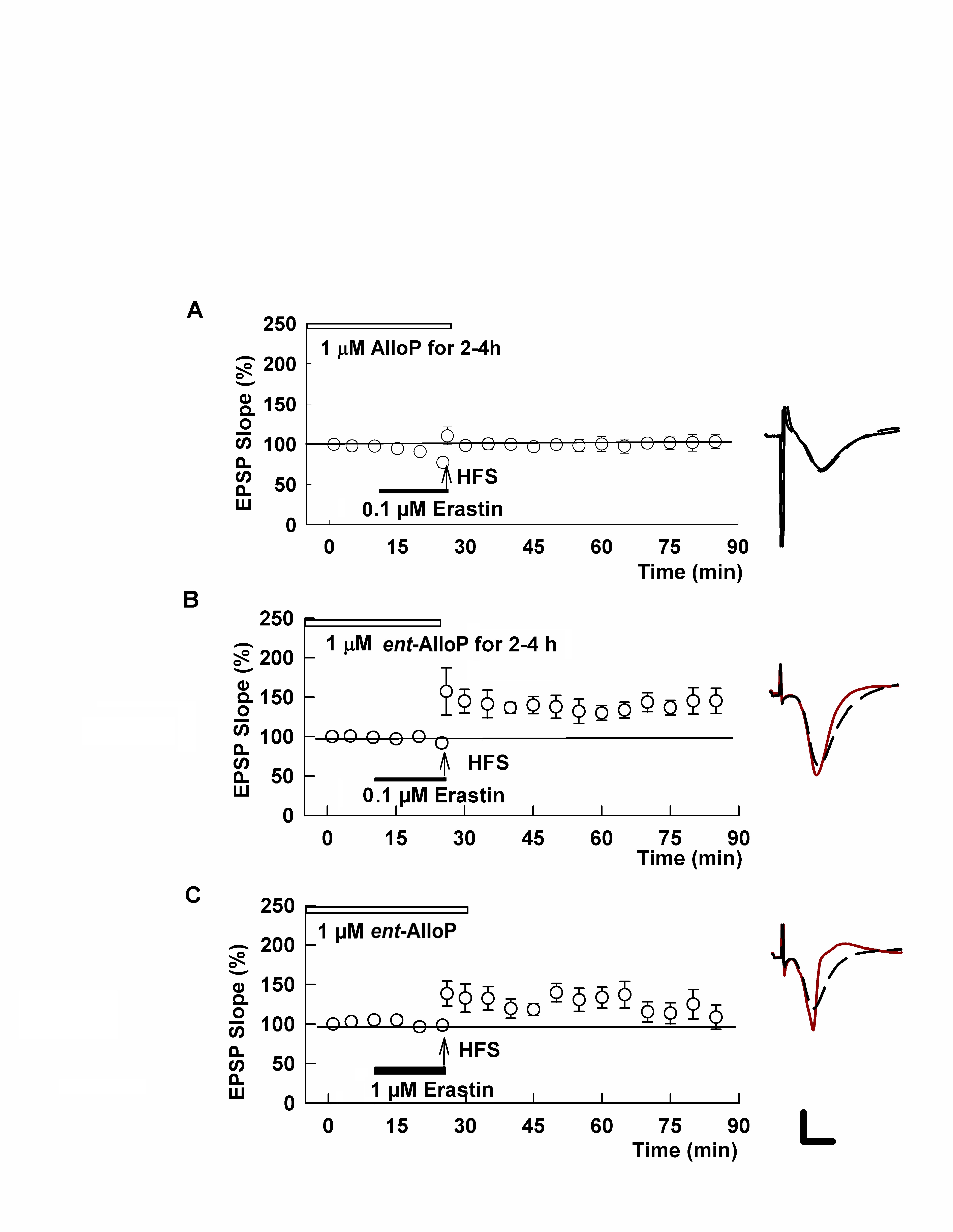

### Extended Figure 5-2

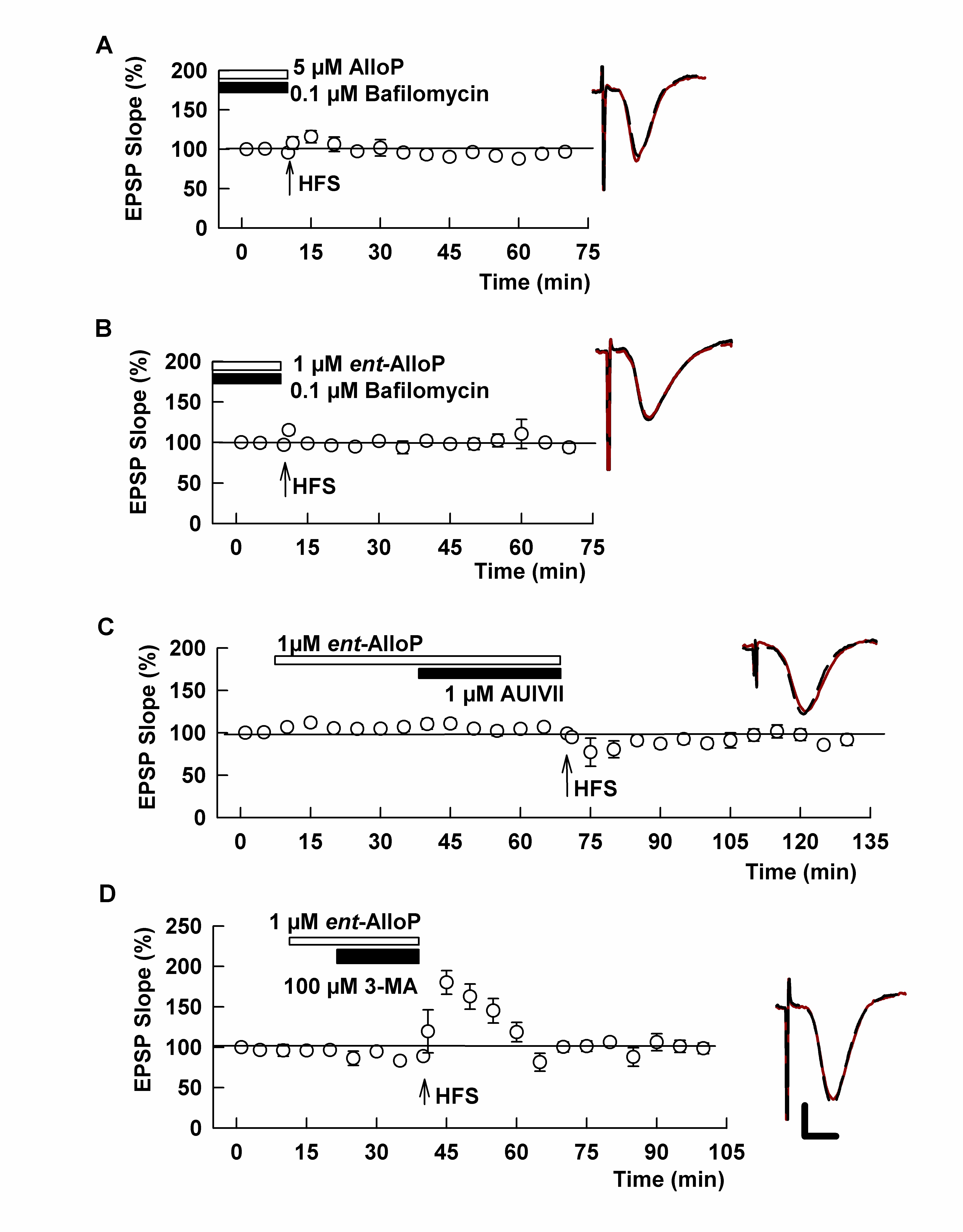
